## Supplementary Material for "Speech prosody enhances the neural processing of syntax"

### Speech Stimuli

During MEG acquisition, participants were presented with 7 different blocks of TED talk recording extracts of various duration, with an average of 509 seconds. The following table shows how the 4 TED talks were split in the experiment.

| *Block* | *File audio* | *Length (s)* | *#Words* |
|---|---|---|---|
| *1* | DanielKahneman 2010_part1 | 516 | 1358 |
| *2* | DanielKahneman 2010_part2 | 488 | 1304 |
| *3* | JamesCameron 2010_part1 | 474 | 1397 |
| *4* | JamesCameron 2010_part2 | 519 | 1573 |
| *5* | JaneMcGonigal 2010_part1 | 549 | 1775 |
| *6* | JaneMcGonigal 2010_part2 | 631 | 2058 |
| *7* | TomWujec 2010 (entire talk) | 387 | 1124 |

### Prevalence score and null distribution assessment of cortical decoding of MEG data

We computed the prevalence values for the MVPA analysis to assess the within subject neural decodability of the three conditions in the word pre- and post-offset time windows, following a Bayesian approach[1]. The within participant significance tests were computed with a permutation test of the AUC values. Each permutation was extracted by shuffling the original labels across all the three conditions 1000 times. For the pre-offset MVPA of the coherent condition, with mean AUC of 0.52 (pVal < 0.001), we found a population prevalence Bayesian maximum a posteriori estimate (MAP) of 0.71 (95% HPDI :[0.42 0.91]). The incoherent condition, with mean AUC of 0.5 (pVal = 0.207), showed a population prevalence MAP of 0.14 (95% HPDI :[0.00 0.41]). For the neutral condition, with mean AUC of 0.51 (pVal < 0.001), we found a population prevalence MAP of 0.43 (95% HPDI :[0.17

0.70])) (Fig. 1A). These results suggest that the coherent condition has the best within participants decoding with a good population prevalence of true positive. This is demonstrated by a MAP estimate greater than 0.5 (majority threshold).

Better trends across conditions were found in the post-offset time window, although with higher MAP values (Fig. 1B). Again the coherent condition shows the highest population prevalence, with mean AUC of 0.53 (pVal < 0.001) and a population prevalence MAP of 1.00 (95% HPDI :[0.77 1.00]). For the incoherent condition, with mean AUC of 0.51 (pVal = 0.034), we found a population prevalence MAP of 0.62 (95% HPDI :[0.33 0.85]). Finally, for the neutral condition, with mean AUC of 0.52 (pVal = 0.001), we found a population prevalence MAP of 0.90 (95% HPDI :[0.64 0.99]) . These values of prevalence score thus confirm the robustness of the decoding analysis also in the post-offset window, where all the conditions showed MAP estimate greater than 0.5.

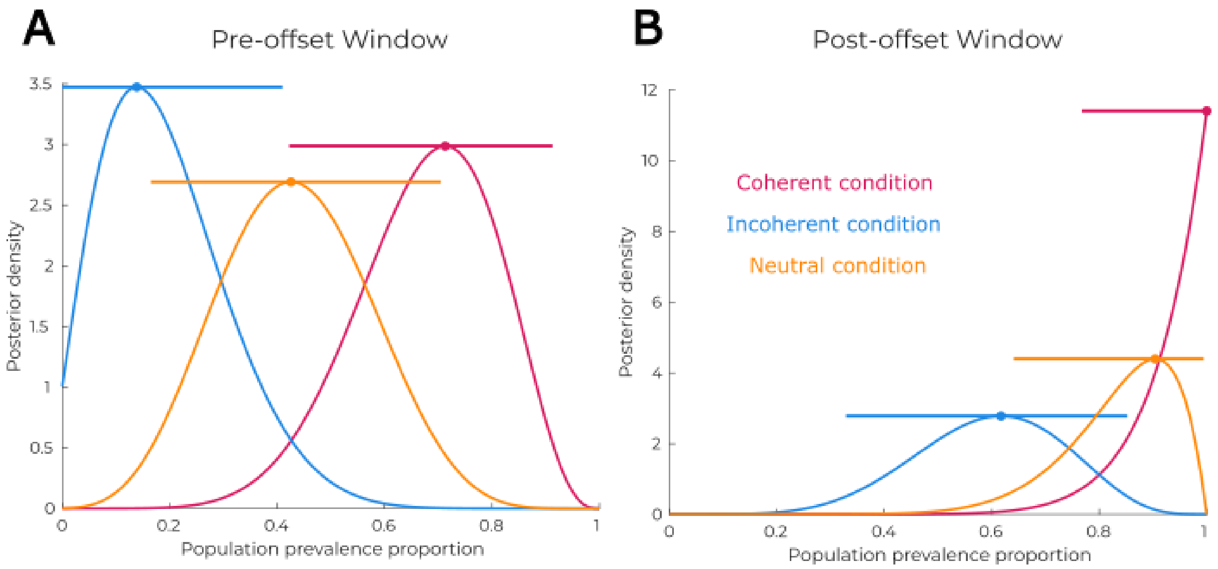


*Figure 1: Posterior distribution of prevalence scores of the three conditions in the two time windows before (panel A) and after token offset (panel B).*

The permutation test also allowed us to assess if the assumption of a chance level AUC of 0.5 was valid. None of the within-participants AUC scores, across conditions and time-windows, showed that the chance level of AUC = 0.5 was different from the null distribution given by the permutation. Figure 2 shows the histograms of the results of the 1000 permutation of the first 5 participants for each condition.

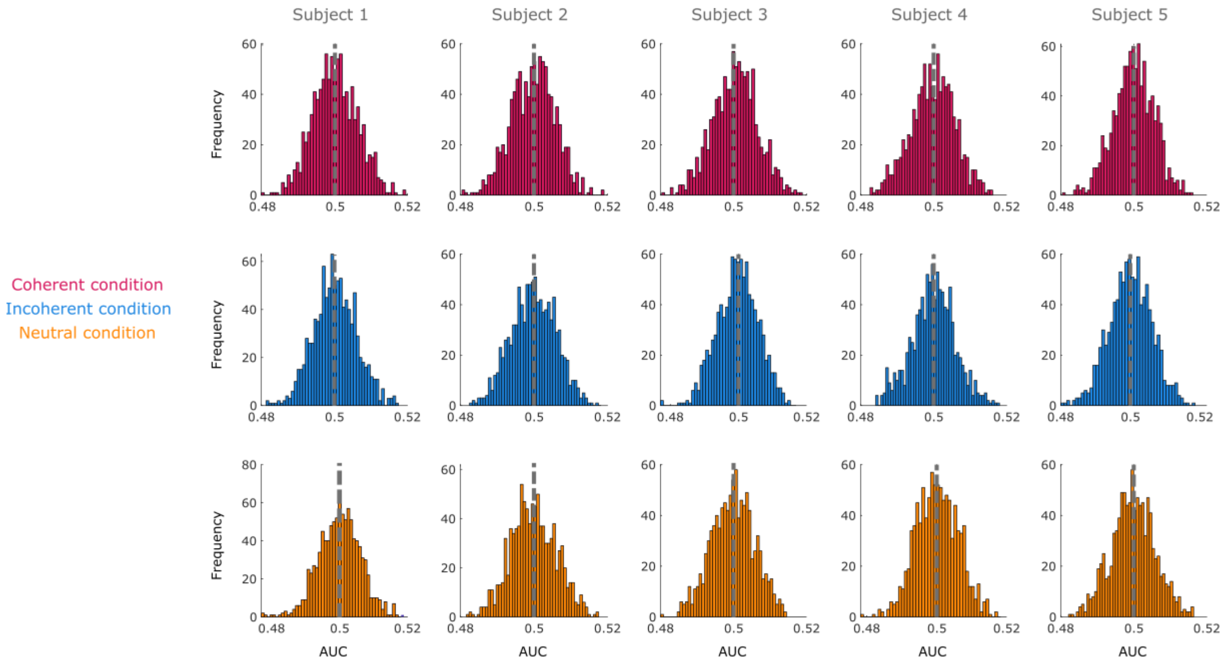


*Figure 2: Histograms of the distribution around chance level of 0.5 for the AUC. Columns correspond to 5 examples of subjects while rows are the results across the three different conditions. All distribution within subjects seems to follow a normal distribution*

### Test set results across the four binarized syntax by prosody conditions

We further unpacked the decoding results in the 4 different classes based on the presence of closing phrase boundaries (cPhBound) and on the level of prosodic boundary (PB) strength (Fig. 3).

The permutation test in the low prosodic boundary strength and cPhBound condition as well as that in the high prosodic boundary strength and no cPhBound condition did not show any significant cluster. Decoding from words with low prosodic boundary strength and no cPhBound revealed multiple clusters after word offset (Cluster 1: [0.056, 0.103]s, pVal = 0.036; Cluster 2: [0.177, 0.223]s, pVal = 0.021; Cluster 3: [0.391, 0.438]s, pVal = 0.030). Similarly, the condition with strong prosodic boundary strength and phrase boundary showed significant temporal decoding clusters that spanned across both pre- and post-word offsets, with a peak in the first 200ms post word-offset (Cluster 1: [-0.065, 0.056]s, pVal = 0.006; Cluster 2: [0.069, 0.210]s, pVal = 0.004; Cluster 3: [0.223, 0.277]s, pVal = 0.021; Cluster 4: [0.297, 0.357]s, pVal = 0.021). These latter two significant test sets together correspond to the coherent condition, and therefore converge with the results presented in the paper.

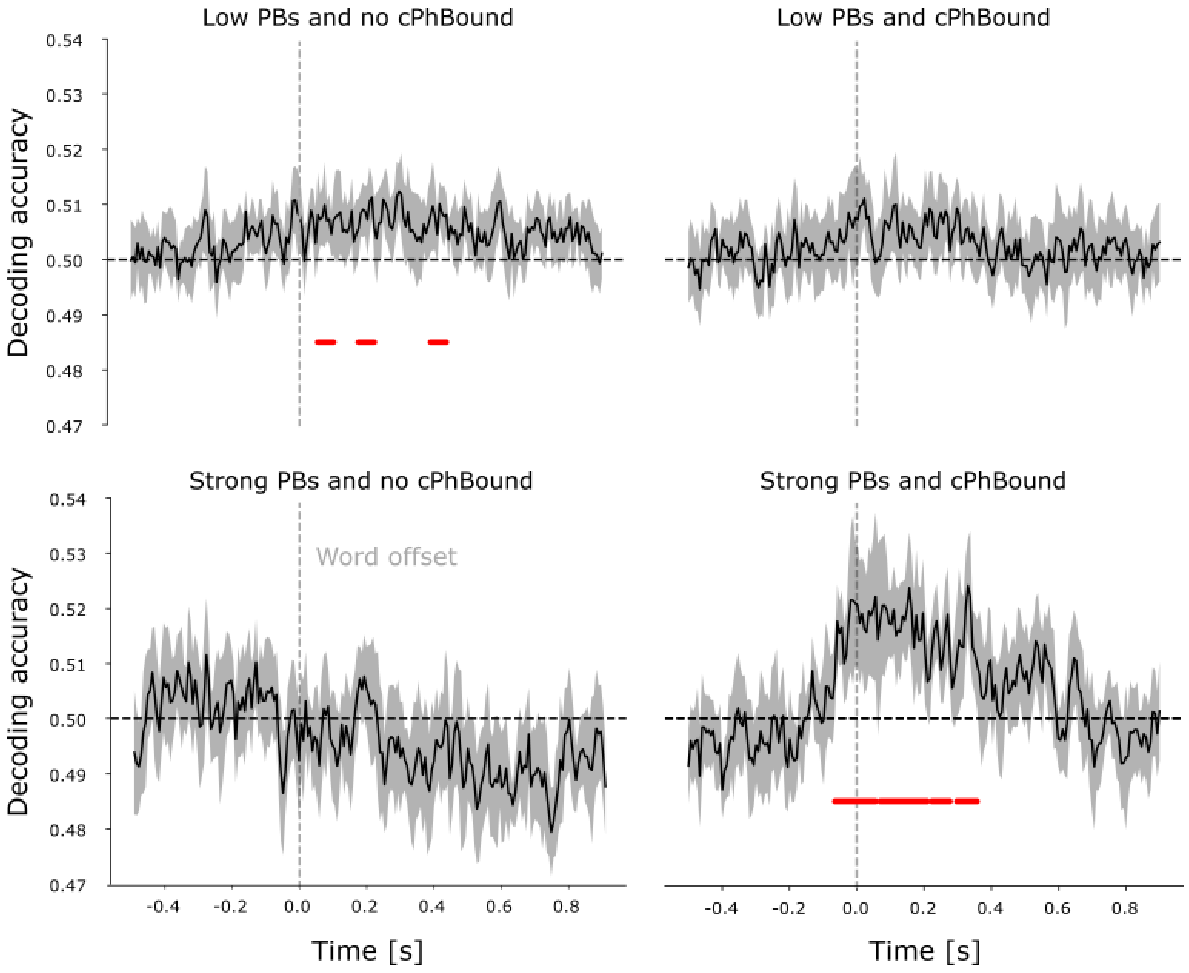


*Figure 3: Generalization sets according to the four different classes of category combinations (low and high prosodic boundary strength vs presence or not of closing phrase boundaries). Red lines show significant clusters (p<0.05).*
